# Low STING expression in a transplantable Kras^G12D^/P53^ko^ lung cancer model contributes to SiglecF^+^ neutrophil and CD103^+^Treg accumulation in tumors

**DOI:** 10.1101/2021.01.04.425311

**Authors:** Laurent Gros, Chiara Ursino, Julie Constanzo, Nadine Zangger, Etienne Meylan, Nathalie Bonnefoy, Julien Faget

## Abstract

Lung cancer is the leading cause of mortality by cancer worldwide. Non-small cell lung cancer is the most common type of lung cancer and mutations in the *KRAS* gene are frequently found in this pathology. While immune checkpoint inhibitors are providing new hope for lung cancer care, only a subset of patients show durable benefit from these new therapies designed to drive an efficient anti-tumor immune response. Hence, it is crucial to better understand the mechanisms through which the tumor immune microenvironment is established in lung tumors. Using bioinformatics, we observed that high expression of the STimulator of INterferon Gene (STING) associates with a longer overall survival specifically in KRAS mutant cancer patients. In lung cancer cell lines, STING expression is linked to interferon response and epithelial-to-mesenchymal transition. Because STING activation in immune cells of the tumor microenvironment using specific agonists is an emerging strategy to trigger an anti-tumor immune response, we took advantage of two transplantable models of *Kras* driven lung cancer, expressing high or low levels of STING, to investigate the function of STING directly in cancer cells *in vivo*. We observed that high-STING expression and constitutive STING signaling were critical for transplanted tumor formation rather than playing a major role in tumor immunogenicity. Besides, low-STING expression in cancer cells is associated with an immunosuppressive tumor microenvironment characterized by the accumulation of tumor promoting SiglecF^+^ neutrophils and CD103^+^ regulatory T cells. In that model, knocking out STING increased the early response to anti-PD1 treatment. We conclude that low-STING expression in cancer cells might confer them an independence from pro-inflammatory signals and a greater immunosuppressive capability and aggressiveness.

## Introduction

Immune checkpoint blockade is the latest revolution in the care of patients with lung cancer. The anti-programmed cell death receptor or ligand-1 (PD1/PDL1) antagonist antibodies, which work by restoring the anti-tumor activity of exhausted CD8^+^ T cells^1,2^, are giving promising results, however, only 22% of all lung cancer patients respond to anti-PD1/PDL1 immunotherapy^3^. For non-small cell lung cancer (NSCLC) patients, the Checkmate-052, KEYNOTE-001 and KEYNOTE-010 clinical trials indicated that patients with a PDL1 positive tumor cell proportion score superior or equal to 50% are more likely to respond to pembrolizumab (anti-PD1). This finding provided the first mechanism-driven biomarker and revealed that only a 30.8% of lung cancer patients (KEYNOTE-024 clinical trial) were indeed eligible for nivolumab/pembrolizumab therapy^4,5,6^. Together with PDL1 expression, tumor mutation burden (TMB) and total CD8 T cell infiltration are used as predictors of immunotherapy success. Despite the fact that lung cancer is considered as immunogenic, tumors presenting a high level of infiltration by immune cells, developing strategies for modifying the composition of the tumor immune microenvironment could be a corner stone to enlarge the spectrum of patients that could benefit from immunotherapy. Hence, it is critical to better understand the interplay between cancer cell properties and the tumor microenvironment.

Lung tumor cell intrinsic signaling pathway alterations can play a major role on the response to immune checkpoint blockade^7^. In lung cancer, the inactivation of PTEN^8^ or LKB1 (also known as STK11)^9^ tumor suppressor associates with resistance to anti-PD1 treatment. Although described in a small number of patients, the alteration of interferon-γ signaling and antigen presentation, in the cancer cells, also constitute escape mechanisms to immune checkpoint blockade^8,10^. On the other hand, the impact of tumor cell innate immune signaling is not well understood.

The STimulator of INterferon Gene (STING) pathway recently emerged as an important player in the establishment of an effective anti-tumor immune response. Canonical STING signaling is induced by the conversion of extranuclear DNA into cyclic Guanosine(2’,5’)phosphate-Adenosine(3’,5’)phosphate (2’3’-cGAMP)^11^, the natural ligand of STING, by the cyclic GMP-AMP synthase (cGAS)^12^. This drives the phosphorylation of interferon regulatory factor-3 (IRF3) by Tank-binding kinase-1 (TBK1), ultimately leading to the transcription of interferon-stimulated genes (ISG). The canonical STING pathway is a potent inducer of *CXCL10, CCL5, IRF9* and interferon-alpha, beta and/or gamma (IFNα/β and γ) in multiple cell types^13^.

Through its pivotal role as innate immune sensor involved in antiviral immunity, triggering STING activation in the tumor immune compartment following radiation therapy^14^ or using various specific STING ligands was shown to reduce tumor growth of CT26, B16 and 4T1^15,16^ cancer models. STING agonist also increases anti-PDL1 efficacy in a tolerized model of breast cancer^17^ and when used as adjuvant for anti-cancer vaccine therapy combined with PD1 blockade in poorly immunogenic cancer models^18^. Because an increasing number of publications reported that STING activation in tumor-infiltrated immune cells might favor immune-checkpoint blockade efficacy, several STING agonists are currently under evaluation in clinical settings^19^. However, the function of STING in the tumor epithelial cells remains poorly understood.

In NSCLC, LKB1 loss in cancer cells leads to STING gene (*TMEM173*) epigenetic silencing and therefore associates with insensitivity to extranuclear DNA sensing, avoiding induction of IRF3 target genes in the cancer cell and facilitating immune escape^20^. On the contrary, impairment of the DNA damage response, through the blockade of CHK1 and PARP, was recently shown to drive extranuclear DNA accumulation in a model of small cell lung cancer. This accumulation led to cGAS-dependent canonical STING signaling in the cancer cells and sensitized lesions to anti-PD1 treatment^21^. Following the same vein of research, low doses of carboplatin administration was shown to activate STING signaling in cancer cells of the Lewis Lung Carcinoma model (LLC); this contributed to increase the efficacy of anti-PD1 antibody^22^.

However, a cGAS-independent alternative STING signaling upon etoposide-induced DNA damage was shown to drive NF-kB activation independently of TBK1/ IRF3-mediated ISG expression in keratinocytes^23^. Furthermore, both NF-kB^24^ activation and a CCL5 autocrine circuit^25^ (one important target of STING signaling) are required in Kras induced mouse models of lung cancer.

We investigated here the impact of endogenous STING expression in two Kras^LSL-G12D/WT^; p53^fl/fl^ (KP) mouse-derived transplantable lung cancer models. One was displaying high-STING expression and showed a strong immunogenicity while the other was characterized by low-STING expression and demonstrated high immunosuppressive capability. In this manuscript, we report that, while it might modestly contribute to anti-tumor immunity in our highly immunogenic lung cancer model, STING plays a major role during transplanted tumor initiation, and low-STING expression contributes to the accumulation of SiglecF^+^ neutrophils and CD103^+^ regulatory T cells (Treg) in immunosuppressive tumors.

## Results

### High STING expression in tumor correlates with longer survival in KRAS-mutant NSCLC patients

We explored the cancer genome atlas (TCGA) database to determine whether STING/TMEM173 expression is linked with the survival of patients with lung adenocarcinoma, the main NSCLC subtype. While STING expression is not linked with survival in this cohort of 198 patients when considered as a whole, these analyses demonstrated that higher STING expression (based on a median split) is positively associated with a longer survival among NSCLC patients harboring a mutation in the *KRAS* gene (60 patients) (figure 1A). STING being ubiquitously expressed and at various levels across cell types, these observations do not infer on the sole impact of cancer-cell STING expression on patient survival as TCGA transcriptomic profiles are also influenced by gene expression from non-tumor cells, including immune cells. We then performed gene-set enrichment analyses^26^ on lung cancer cell lines from the cancer cell line encyclopedia (CCLE) database and observed that STING expression is positively associated with gene-sets of the HALLMARK collection from mSigDB^27^ identified as IFNγ, IFNα response and epithelial-to-mesenchymal transition (figure 1B). Consistently with enhanced interferon signature^10^, increased STING expression across cell lines positively associates with *PDL1* gene transcription in both *KRAS* mutant and wild type cells (figure 1C). Because there are several evidences linking interferon signaling and PDL1 expression^10^, these observations suggest that cancer-cell STING expression associates with an immune microenvironment prone to the establishment of an efficient anti-tumor immune response upon anti-PD1 treatment. With the aim of identifying a relevant model to study STING function in cancer cells during anti-PD1 treatment, we analyzed STING expression in four different lung cancer cell lines derived from *Kras*^LSL-G12D/WT^;*p53*^fl/fl^ (KP) mice and designated as M8, SV2, Clem42 and Clem2. Strikingly, STING expression showed high variability across these different cell lines. STING protein was undetectable in Clem42 cells, M8 and SV2 cells showed low expression while Clem2 cells displayed high STING expression comparable to the level observed in mouse embryonic fibroblasts (MEF) (figure 1D).

**Figure 1:**
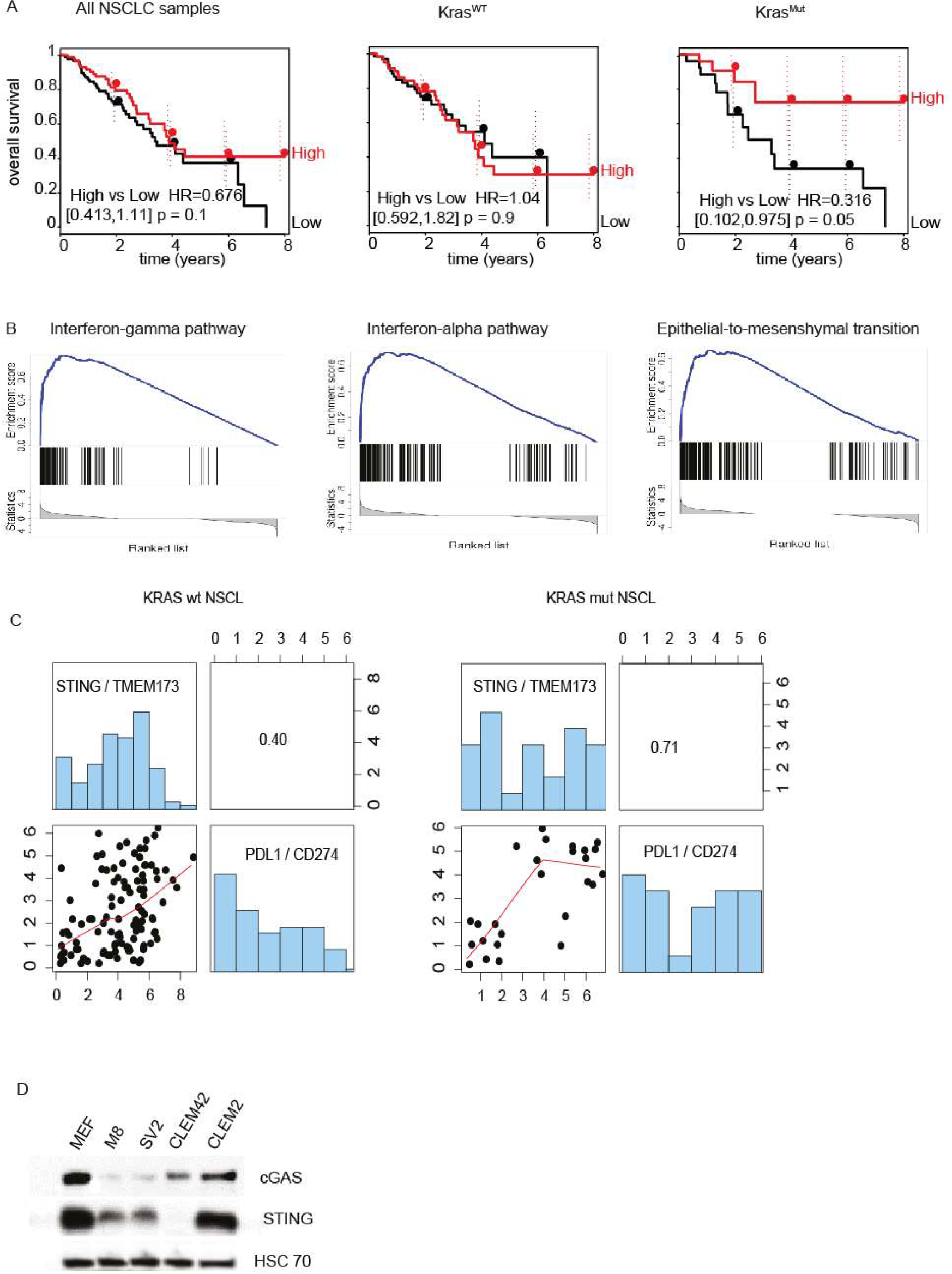
High STING expression associates with KRAS-mutant NSCLC survival. A) Kaplan–Meier curves for overall survival, hazard ratios (HRs), confidence intervals, and *p* values of pairwise differences between groups with high or low *STING/Tmem173* expression in The Cancer Genome Atlas (TCGA) lung adenocarcinoma data sets, Left: all samples (198 patients), middle *KRAS-WT* samples (138) and right: *KRAS-mutant* samples (60 samples). Differences between curves were assessed by using the log rank test (Mantel-Cox). B) Differential gene set enrichment between STING-high and STING-low lung cancer cell lines based on a medial split across samples of the cancer cell line encyclopedia (CCLE) dataset. Graphics depicts enrichments of the INFγ, IFNα response and epithelial-to-mesenchymal transition gene sets from the mSigDB HALLMARK collection. C) Graphics shows correlation between STING/TMEM173 expression and PDL1/CD274 in KRAS-WT and KRAS-mutant cell lines of the CCLE dataset. D) Western-blot showing cGAS, STING and HSC70 protein expression in MEF, M8, SV2, Clem42 and Clem2 cells.

### Lymphocyte-neutrophil infiltration and sensitivity to PD1 blockade differentiates Clem2 and M8 tumors

To compare the ability to form tumors and the tumor microenvironment of STING-high and STING-low cancer cells, we performed orthotopic transplantation of 5×10^4^ STING-high Clem2 and STING-low M8 cells in syngeneic C57BL/6J mice through tail-vein injection. Computed tomography revealed that STING-high Clem2 cells have a lower ability to colonize the mouse lung when compared to STING-low M8 cells (figure 2A). From 7 mice, only 6 died from Clem2 cell transplantation after 18 weeks, while M8 cell transplantation led to the death of the all the engrafted mice within 7 weeks (figure 2B). Trucount flow cytometry at end-point (when mice displayed important breathing defaults and/or body-weight loss superior to 15%) revealed a higher quantity of immune cells per mg of lung tissue in STING-high Clem2 cell-transplanted mice (figure 2C). Furthermore, spleen weight and number of leukocytes per ml of blood were higher in Clem2 than in M8 transplanted mice (supplementary figure 1A and B) suggesting that a systemic alteration of immune cell homeostasis occurs consequently to Clem2 but not to M8 cell transplantation. Deeper analyses of the myeloid compartment showed that lungs of M8 transplanted mice displayed a significantly higher proportion of SiglecF^+^ tumor-promoting neutrophils^28,29^ when compared to Clem2 (figure 2D and E). Conversely, CD8^+^ T cell proportion was lower in the lungs of M8 than in Clem2 transplanted mice while the proportion of CD4^+^ T lymphocytes among immune cells was similar (figure 2F). Hence, these observations demonstrated that orthotopic transplantation of Clem2 cells led to a stronger immune cell infiltration containing lower SiglecF^+^ neutrophil and higher CD8^+^ T lymphocyte proportions in the lung when compared to the M8 model. To evaluate the robustness of this immune signature we decided to analyze the tumor growth and immune landscape of Clem2 and M8 tumors after sub-cutaneous injection. We performed preliminary experiments (not shown) and determined that transplanting 3×10^5^ Clem2 and 1.5×10^5^ M8 cells provided comparable growth rates (figure 2G). Interestingly, subcutaneous tumors showed an immune signature exacerbating the differences already observed in the mouse lungs from the orthotopic model. Clem2 tumors presented a very low proportion of total neutrophils and of SiglecF^+^ neutrophils and a high proportion of CD8^+^ T lymphocytes when compared to M8 tumors (figure 2H and I). In line with our previous observations, Clem2 tumors showed a high CD8^+^ T cell to regulatory T cell (Treg) ratio (figure 2J) and Trucount flow cytometry demonstrated that the quantity of CD8^+^ T lymphocytes was higher and the quantity of neutrophils was lower in Clem2 than in M8 tumors (figure 2K and supplementary figure 1B). These major differences between Clem2 and M8 tumors prompted us to compare their sensitivity to anti-PD1 treatment. Strikingly, Clem2 tumors showed complete response to anti-PD1 antibody treatment while anti-PD1 treatment had no impact on M8 tumor growth (figure 2L).

**Figure 2:**
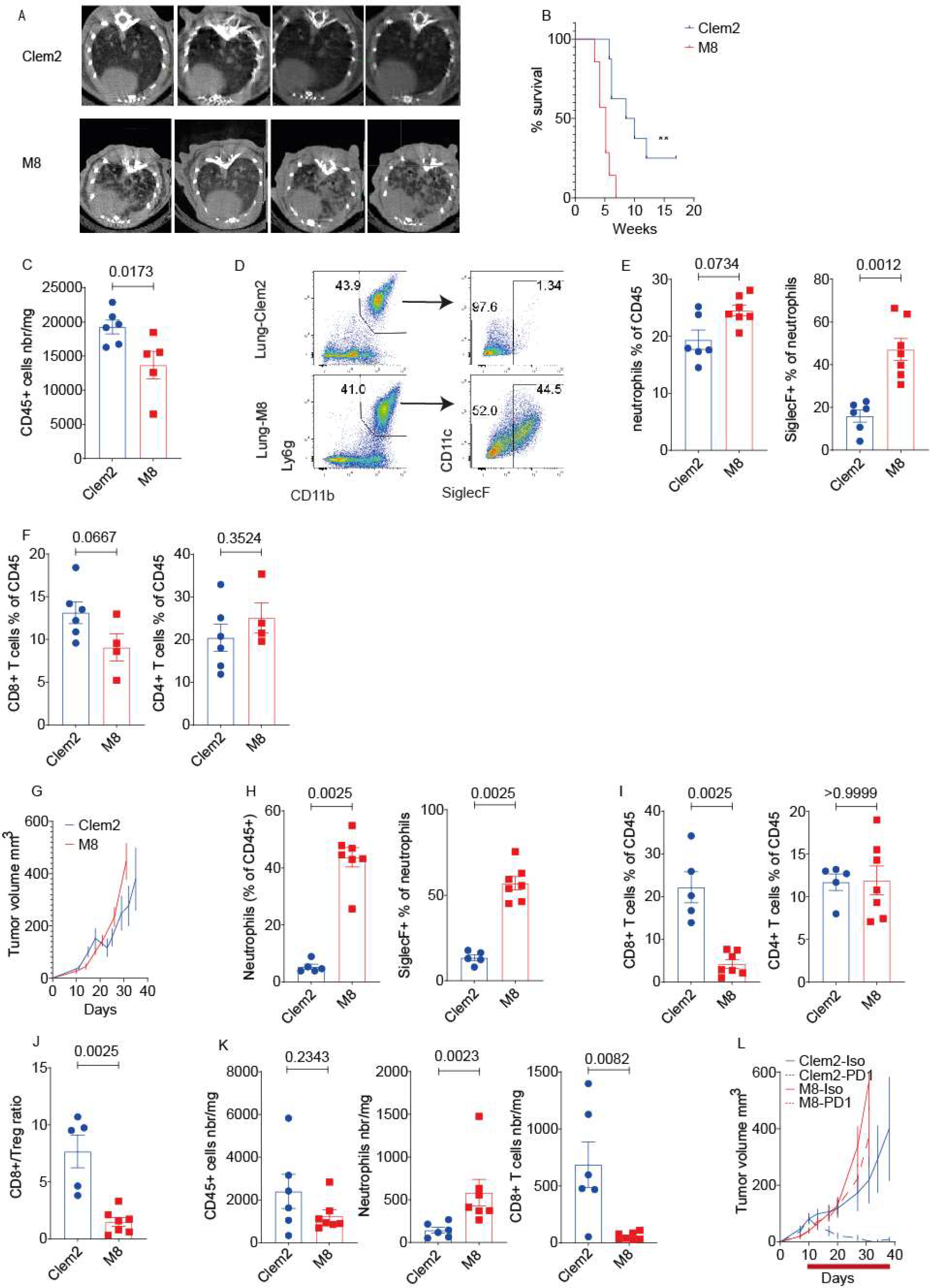
Clem2 and M8 cells forms tumors displaying opposite immune profile and sensitivity to anti-PD1 treatment. A) Micro-computed tomography showing tumor burden at week 5 post tail vein injection of 50×10^3^ Clem2 (top) and M8 (bottom) cells in syngeneic C57BL/6J male mice (n=4 per group). B) Kaplan–Meier curves showing mice overall survival in the same experiment than in A. (n=7 mice per group, statistic was obtained using Wilcoxon test, ** p<0.01). C) Histogram shows the number of immune cells per mg of lung at end-point from Clem2 (n=6) and M8 (n=5) mice accessed by Trucount flow cytometry. D) Representative flow cytometry plot showing gating trategy used to identify SiglecF^+^ neutrophils among viable, linage negative (CD3, CD4, CD8, CD19, NKp46). CD45^+^ immune cells in the mouse luna in Clem2 (top) and M8 (bottom) cells transplanted mice. E) Histograms show the proportion of Ly6G^+^CD11b^+^ neutrophils among CD45^+^ immune cells (left) and the proportion of SiglecF^+^ neutrophils among neutrophils (right) identified as in D in the lung of Clem2 (n=6) and M8 (n=7) transplanted mice. F) Histograms show the proportion of CD3^+^CD8^+^ (left) and CD3^+^CD4^+^ (right) T lymphocytes among viable CD45^+^ immune cells in the same experiment than in E. Clem2 n=6, M8 n=4. G) Curves depict Clem2 and M8 subcutaneous (sc)-tumor growth following the transplantation of 3×10^5^ or 1.5×10^5^ cells respectively, n=7 mice per group. H-J) Histograms show the proportion of H) Ly6G^+^CD11b^+^ neutrophils among CD45^+^ immune cells (left) and the proportion of SiglecF^+^ neutrophils among neutrophils (right), of I) CD3^+^CD8^+^ (left) and CD3^+^CD4^+^ (right) T lymphocytes among viable CD45^+^ immune cells in Clem2 and M8 sc-tumors and of J) the CD8+ T lymphocytes over Treg ratio. n=5 Clem2 and n=7 M8 tumors. K) Histograms depicts numbers of the indicated immune cells population per mg of tumor tissue accessed by true-count flow cytometry. Clem2 n=6, M8 n=7. C, E, F, H-K) numbers indicate p-values obtained from Mann Whitney test: L) Curves shows sc-tumor growth in Clem2 (blue) and M8 (red) syngeneic mice treated with 150μg of control antibody (filled line) or anti-PD1 antibody (dashed line) twice a week, starting at day 10 post-transplantation and up to the end of the experiment. N=7 mice per group.

To conclude, STING-high Clem2 cancer cells form tumors highly infiltrated by CD8^+^ T lymphocytes and are characterized by a strong sensitivity to PD1 blockade; conversely, STING-low M8 tumors are resistant to anti-PD1 antibody treatment and generate lesions highly infiltrated by SiglecF^+^ tumor promoting neutrophils.

### STING-IRF3 signaling is constitutively activated in Clem2 cells but fails to induce significant *Ifnb* transcription

Before the evaluation of the link between STING expression and the tumor immune microenvironment in mice, we explored the functionality of the STING pathway in these two cell lines *in vitro*. Because STING signaling drives IRF3 target gene expression, we dissected the STING/IRF3 pathway in MEFs and lung cancer cells by transducing MEFs, Clem2 and M8 cells with a lentiviral vector carrying a dominant negative form of IRF3 (ΔN-IRF3), downstream to tetracycline-inducible enhancer sequences. Hence, doxycycline treatment was expected to induce ΔN-IRF3, leading to IRF3-dependent gene inhibition. We first validated the efficient induction of ΔN-IRF3 after 72 hours of doxycycline exposure (supplementary figure 2A). Next, we treated the stable cells with the specific STING ligand, 2’3’-cGAMP for 4 hours, with or without prior ΔN-IRF3 induction and monitored *Cxcl10, Ccl5*, and *Infb* gene transcription by real-time PCR. We observed that 2’3’-cGAMP treatment led to a 31-, 1.8- and 13.9-fold induction of *Cxcl10* transcription in MEFs, Clem2 and M8 cells respectively. STING-mediated induction of *Cxcl10* was partially (about 30%) inhibited by ΔN-IRF3 expression in MEFs and M8 cells upon 2’3’-cGAMP treatment, and in the control and 2’3’-cGAMP treated conditions for Clem2 cells that have high basal *Cxcl10* levels when compared to M8 cells (figure 3A).

**Figure 3:**
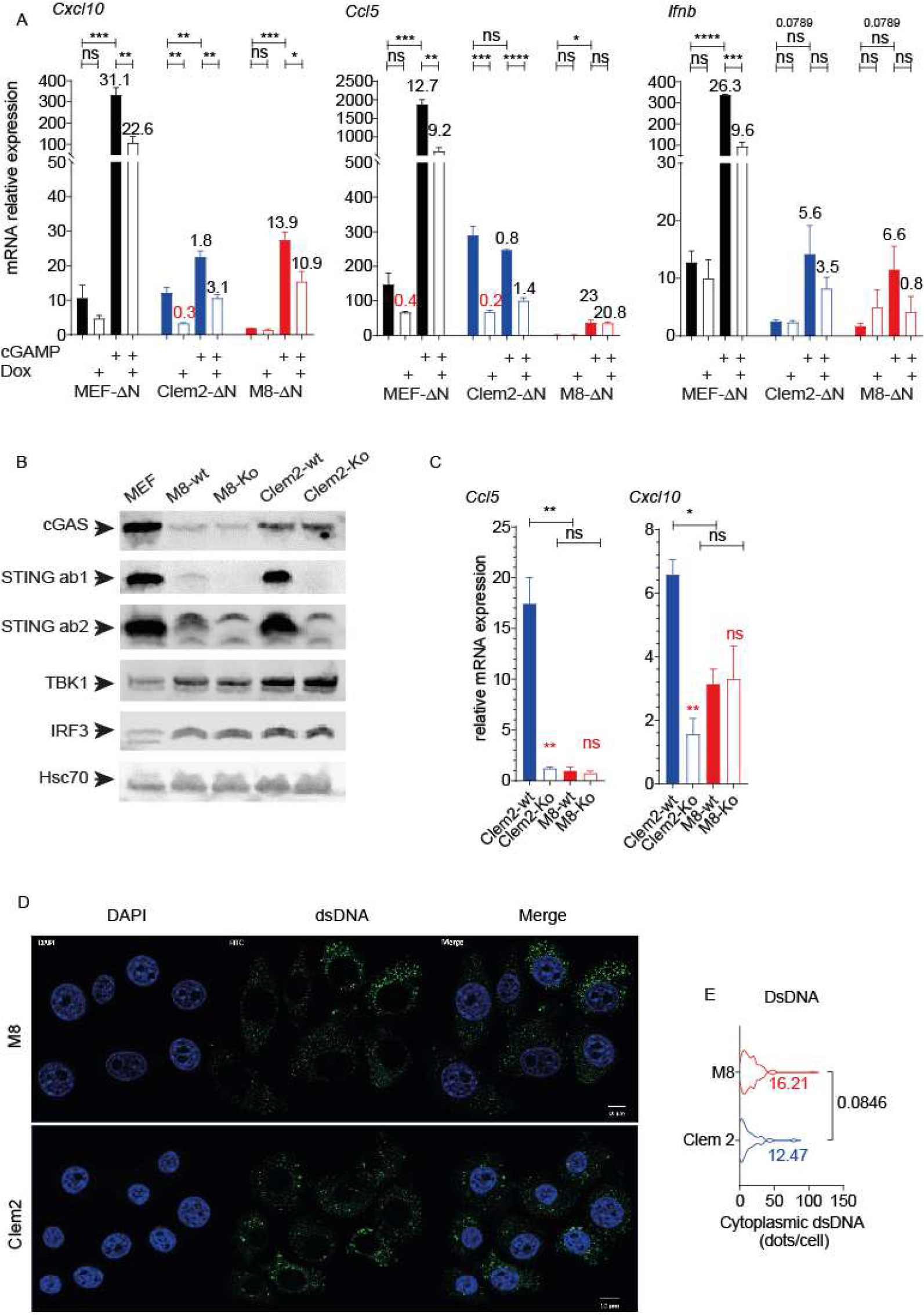
STING signaling is constitutively active in Clem2 cells. A) Histograms show relative mRNA expression of the indicated gene (left:Cxcl10, middle Ccl5, right *Ifnb*) in MEF, Clem2 and M8 cells expressing (empty bars) or not (filled bars) ΔN-IRF3 (Dox: pre-treated with doxycycline for 72 hours) then treated with lipofectamine containing medium of lipofectamine plus 10μg/ml of 2’3’cGAMP during 4 hours before RNA extraction. Relative mRNA expression was calculated from ΔCT respectively to *Gapdh* expression. Black numbers indicate fold change in mRNA expression relative to the respective control without 2’3’cGAMP. Red numbers indicate the fold change between Dox versus no-Dox in absence of 2’3’cGAMP. B) Western blot showing cGAS, STING, TBK1, IRF3 and HSC70 expression in MEF, M8-wt, M8-Ko, Clem2-wt and Clem2-Ko cells. M8-wt and Clem2-wt were transduced with the same CrispR vectors than M8-Ko and Clem2-Ko but lacking the guild RNA driving STING/Tmem173 deletion. STING ab1 and STING ab2 correspond to the monoclonal anti-STING antibody: Cell Signaling (ref: 13647) and the polyclonal anti-STING antibody: Proteintech (ref 19851-1-AP) respectively. A, C, E) * p<0.05, ** p<0.01, ***p<0,001, Statistics were obtained using student-T test, figures show representative results of one experiment performed in biological triplicate, each of them were reproduced at least twice.

Similarly, 2’3’-cGAMP led to the induction of *Ccl5* in MEFs (12.7-fold) and M8 (23-fold) cells, with the latter showing almost undetectable mRNA levels of *Ccl5* in basal conditions. On the other hand, *Ccl5* expression was not changed in Clem2 cells that already expressed high levels of this chemokine in absence of STING triggering. Furthermore, in MEFs ΔN-IRF3 expression reduced partially but significantly *Ccl5* expression upon 2’3-cGAMP treatment while it did not revert its induction in M8 cells and strongly decreased it in Clem2 cells with or without exogenous STING ligand exposure (figure 3A).

*Ifnb* expression was induced in MEFs (26-fold) during 2’3’-cGAMP treatment in an IRF3-dependent manner as ΔN-IRF3 reduced *Ifnb* induction by more than 50%. In Clem2 and M8 cells, 2’3’-cGAMP treatment led to a 5.6- and 6.6-fold induction of *Ifnb* transcription that, even if not significant, seemed to be partially dependent on IRF3 (figure 3A).

Strikingly, STING/IRF3 signaling was weak when compared to Ccl5, *Cxcl10* and *Ifnb* induction following Toll-Like Receptor 3 (TLR3) triggering with polyI:C. Indeed, polyI:C induced *Cxcl10* by 6228-, 18.5- and 722-fold, *Ccl5* by 1664-, 5.6- and 4714-fold, and *Ifnb* by 17959-, 294- and 3026-fold in MEFs, Clem2 and M8 cells, respectively (supplementary figure 2B). These observations indicate that STING is not a strong inducer of the antiviral immune response when compared to TLR3 and that, even if Clem2 cells express higher basal levels of *Ccl5* and *Cxcl10*, they have a lower ability to upregulate antiviral genes following STING or TLR3 triggering than MEFs and M8 cells. Furthermore, contrasting to M8 cells, the basal expression of *Ccl5* and *Cxcl10* can be inhibited by ΔN-IRF3 in STING-high Clem2 cells, suggestive of a constitutive IRF3 activity.

To determine if high STING expression is responsible for constitutive IRF3 activation in Clem2 cells, we generated STING-knockout (STING-Ko) Clem2 and M8 cells using the CRISPR/Cas9. We validated that the knockout of STING did not alter cGAS, IRF3 and TBK1 expression and observed that M8 cells differ from Clem2 cells not only by their low expression of STING but also of cGAS (figure 3B). Gene expression analyses revealed that basal expression of *Ccl5* and *Cxcl10* was strongly dependent on STING in Clem2 cells. Indeed, STING-Ko Clem2 cells had a decreased expression of these 2 genes down to the levels observed in M8 cells (figure 3C). Because these observations suggested that constitutive STING signaling occurs in Clem2 cells, we performed immunofluorescence staining against double-stranded extranuclear DNA and observed that both Clem2 and M8 cells contain cytoplasmic DNA susceptible of inducing STING signaling (figure 3D and E).

Finally, to determine if basal STING activation could have an impact on lung cancer cell behavior *in vitro*, we performed proliferation, wound healing and clonogenic assays on M8 and Clem2 STING-wt and STING-Ko cells. These experiments showed that while cell proliferation was not impacted and clonogenicity slightly decreased in M8 cells, while STING deletion significantly reduced the migration capability of Clem2 cells (supplementary figure 2C).

Altogether, these results demonstrated that the STING/IRF3 pathway is partially functional in cancer cells, since artificial triggering of STING signaling induces *Ccl5* and *Cxcl10* but fails to augment significantly *Ifnb* transcription. Furthermore, STING-mediated ISG expression appeared extremely weak in MEF and cancer cells, when compared to the transcription profiles obtained following TLR3 activation, while Clem2 cells showed a low ability to upregulate these genes irrespectively of the signaling pathway (STING or TLR3), when compared to MEF and M8 cells. Finally, we observed a constitutive STING/IRF3 signaling in Clem2 cells driving *Cxcl10* and *Ccl5* expression but not in M8 cells, which could be explained by low STING and cGAS expression in the latter.

### STING expression by cancer cells contributes to tumor formation and shows opposite impact on Clem2 and M8 tumor response to anti-PD1 treatment

To evaluate the impact of the high-STING expression observed in Clem2 cells on the tumor immune microenvironment, we transplanted 3×10^5^ STING-wt and STING-Ko Clem2 cells to generate sub-cutaneous tumors in syngeneic mice. Strikingly, STING-Ko Clem2 cells showed a profound reduction in their ability to form tumors *in vivo*. Indeed, only 2 mice over 7 developed tumors in the STING-Ko group while STING-wt Clem2 cells formed tumors in all of the mice (n=7); however, 1 of the 7 tumors in the STING-wt group regressed spontaneously at day 10 post-transplantation (figure 4A). This important observation indicates that STING expression and potentially constitutive signaling that occurs in STING-high Clem2 cells plays a major role in tumor formation in vivo. However, by increasing the number of transplanted cells to 9×10^5^, both, STING-Ko and STING-wt Clem2 cells formed tumors (figure 4B).

**Figure 4:**
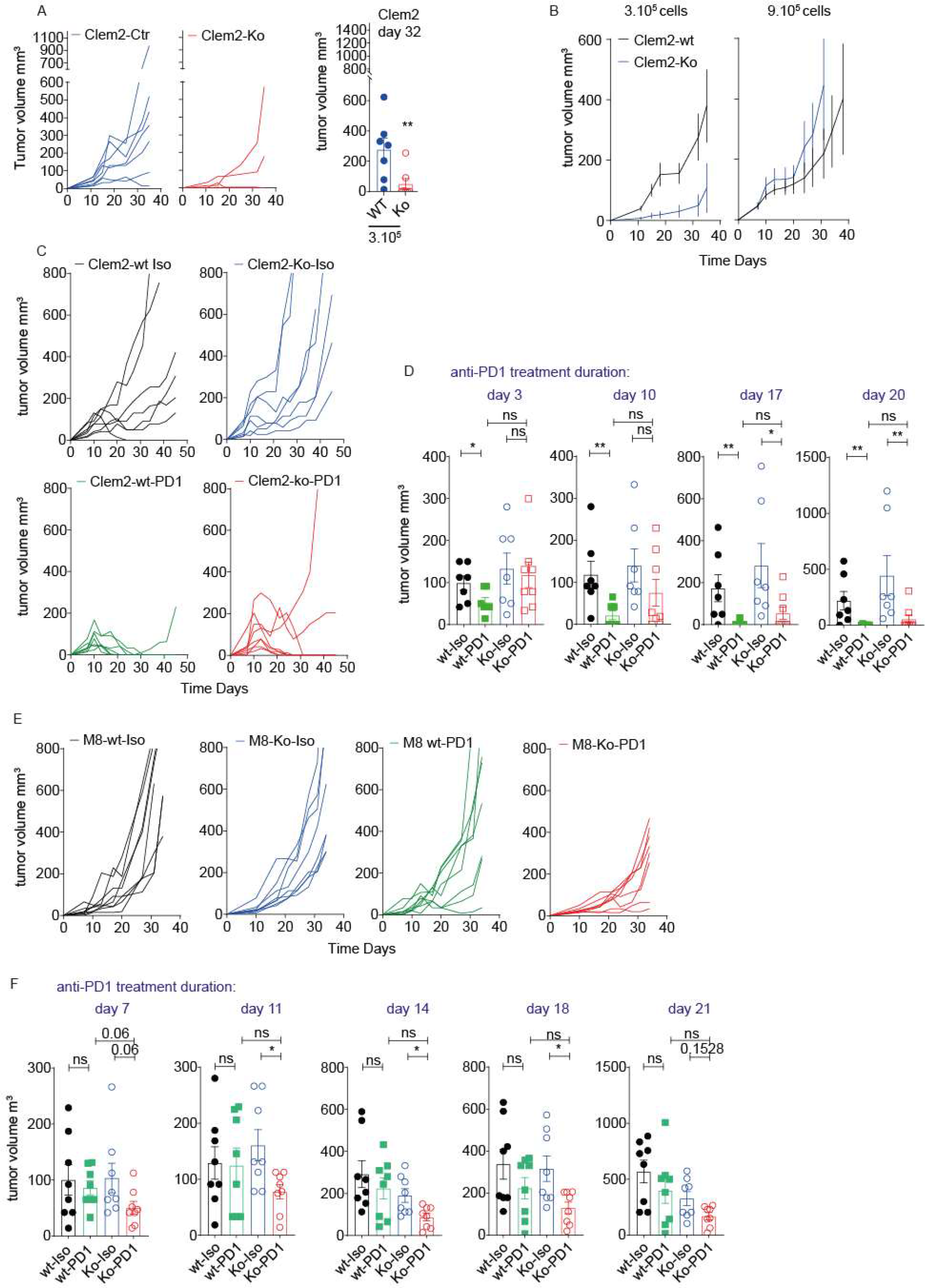
Opposite impact of STING knockout in Clem2 and M8 cells on the early response to anti-PD1. A) Curves indicate individual Clem2-wt (Blue) and Clem2-Ko (Red) sc-tumors growth following the transplantation of 3×10^5^ cells. Histogram depicts individual tumor volumes at day 32 post-transplantation in the same experiment (n=7 mice per group). B) Curves represents average tumor gtowth in Clem2-wt (Black) and Clem2-Ko (Blue) sc-tumors growth after transplantation of 3×10^5^ (left) or 9×10^5^ cells (right) cells (n=7 mice per group). C) Cures shown individual growth of Clem2-wt and Clem2-Ko sc-tumors in mice treated with isotopic control antibody (Celm2-wt: black, n=7; Clem2-Ko: blue, n=7) or anti-PD1 antibody (Clem2-wt: green, n=7; Clem2-Ko: red, n=8). Antibody treatment started at day 10 post-transplantation of 9×10^5^ cancer cells. D) Histogram show individual tumor volume at the indicated time-point from anti-PD1 or isotopic control treatment initiation i n the same experiment than B. E) Cures shown individual growth of M8-wt and M8-Ko sc-tumors in mice treated with isotopic control antibody (M8-wt: black, n=7; M8-Ko: blue, n=7) or anti-PD1 antibody (M8-wt: green, n=7; M8-Ko: red, n=7). Antibody treatment started at day 10 post-transplantation of 1.5×10^5^ cancer cells. F) Histogram show individual tumor volume at the indicated time-point from anti-PD1 or isotopic control treatment initiation in the same experiment than C. A-C and E) * p<0.05, ** p<0.01, ***p<0,001, Statistics were obtained using Mann Whitney test.

In this setting, we observed a high variability of STING-wt Clem2 tumor growth, 2 lesions reached very quickly an exponential growth rate, 4 showed low growth rate over several weeks and 1 demonstrated a complete spontaneous regression starting at day 10 posttransplantation, while in the STING-Ko group, all tumors ultimately reached an exponential growth rate. In other words, the proportion of tumors showing spontaneous regression and slow growth rate over all the experiment duration seemed to be reduced in STING-Ko Clem2 transplanted mice when compared to Clem2 STING-wt (figure 4C). We then treated these mice with anti-PD1 antibody. Four weeks of anti-PD1 treatment led to complete regression of 6 tumors over 7 and 6 tumors over 8 in the STING-wt and STING-Ko Clem2 models, respectively. Interestingly, anti-PD1 showed a very strong effect independently of STING, however we noticed that tumor regression was delayed in the STING-Ko group. Indeed, while anti-PD1 led to a significant tumor volume regression since day 3 post-treatment initiation on Clem2 STING-wt tumors, the effect of anti-PD1 was only visible at day 10 and became significant after 17 days of treatment (figure 4D). Finally, comparative analyses of the tumor immune compartment of Clem2 STING-wt and STING-Ko tumors did not reveal significant differences. However, we observed a trend toward an increase in CD4^+^ T cells and a reduction in CD8^+^ T cells proportions among immune cells in STING-Ko tumors. These variations of lymphocyte proportions associated with a clear trend toward a reduction of the ratio CD8^+^/Treg in STING-Ko Clem2 tumors at endpoint (supplementary figure 3A-C).

We then performed similar experiments using STING-wt and STING-Ko M8 cells. Contrasting to Clem2 cells, STING knockout did not significantly change subcutaneous tumor formation and growth after transplantation of 1.5×10^5^ cells. However, STING-Ko M8 cells displayed a lower ability to colonize the mouse lung, leading to an increased mouse survival after tail vein injection of 5×10^4^ cells (supplementary figure 3D and E). Surprisingly, treatment of mice bearing subcutaneous M8 tumors revealed that STING deletion sensitized the lesions to anti-PD1 (figure 4E). This consisted in a significant delay of tumor growth upon anti-PD1 treatment that was only observed in M8-STING-Ko tumors and lasted from day 7 to day 18 (figure 4F). However, ultimately most tumors became resistant to anti-PD1 treatment.

Altogether, these observations showed that Clem2 cells are dependent on STING as its knockout reduced their ability to form tumors in syngeneic mice. The same phenomenon might occur with a lower intensity in the M8 model as STING-Ko M8 cells were less efficient in colonizing the mouse lung after tail vein injection. Furthermore, we observed a divergent impact of STING knockout on Clem2 and M8 tumors response to anti-PD1. Indeed, while STING knockout decreased the early response to anti-PD1 of Clem2 tumors, it led to a partial but significant sensitization of M8 tumors to this therapy.

### STING-deficient M8 tumors are characterized by lower SiglecF^+^ neutrophil and CD103^+^ Treg infiltration and high proportion of T-bet^+^ CD4^+^ and CD8^+^ T lymphocytes

In accordance with a potential increased immune surveillance and sensitivity to anti-PD1 immunotherapy, quantitative flow cytometry showed a higher quantity of immune cells per mg of tumor tissue in STING-Ko when compared to STING-wt M8 tumors (figure 5A). Furthermore, while the percentages of CD4^+^ and CD8^+^ T cells among the tumor immune infiltrate was not changed, we observed that the proportion of Treg was significantly decreased in STING-Ko M8 tumors (figure 5B). A deeper analysis of Treg cells, indicated that STING knockout in M8 cancer cells associated with a reduction of the proportion of CD103^+^ Tregs (figure 5C). These CD103^+^ Tregs have been described in multiple models of cancer as tumor infiltrating Tregs showing higher immunosuppressive function than their CD103^-^ counterpart^30^.

**Figure 5:**
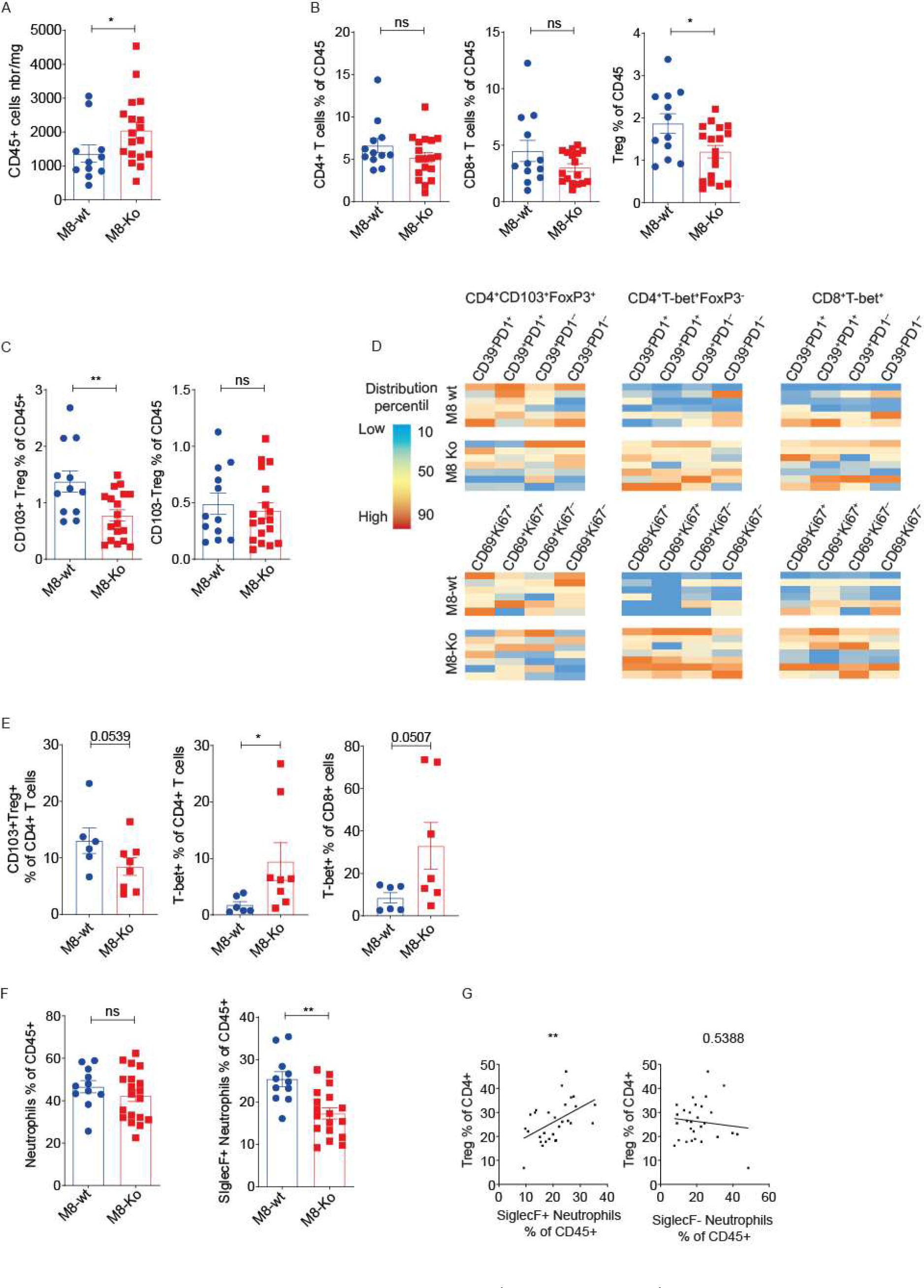
Low-STING expression contributes to CD103^+^Treg and SiglecF^+^ neutrophils recruitment in M8 tumors. A) Histogram shows the number of CD45^+^ cells per mg of tumor tissue in M8-wt (n=11) and M8-KO (n=18) tumors. B) Histograms show percentage of CD3^+^CD4^+^, CD3^+^CD8^+^ T cells and CD3^+^CD4^+^Foxp3^+^ Tregs among viable CD45^+^ immune cells in M8-wt (n=12) and M8-Ko (n=18) sc-tumors. C) Histograms depict the proportion of CD103^+^ (left) and CD103^-^ (right) Treg among viable CD45^+^ in the same samples than B. D) Heat-maps represent the relative proportions of the indicated cell sub-populations among CD45+ cells in M8-wt (top, n=6) and M8-Ko (bottom n=7). The full gating strategy is detailed in supplementary figure 4B and C. E) Histograms show the proportion of CD103^+^ Treg (left), T-bet^+^ (middle) cells among CD4^+^ T lymphocytes and T-bet^+^ cells among CD8^+^ lymphocytes in M8-wt (n=6) and M8-Ko (n=8 and 7). F) Histograms represent the proportion of total neutrophils (left) and SiglecF^+^ neutrophils (right) among viable immune cells identified as detailed in supplementary figure 4D in M8-wt (n=12) and M8-Ko (n=17) E-C and F) * p<0.05, ** p<0.01, Statistics were obtained using Mann Whitney test. G) Scatter plot shows the correlation between the proportions of SiglecF^+^ (left) or SiglecF^-^ (right) neutrophils among immune cells and the proportion of Treg among CD4^+^ T cells across all samples. ** p<0.01 obtained from Pearson test, n=29.

Consistently, further analyses revealed that CD103^+^ Treg cells present in M8 tumors displayed high CD39 expression when compared to CD8^+^ and other CD4^+^ T cell sub-populations and were also characterized by a lower proliferation rate when compared to Foxp3^+^ CD103^-^ Tregs (supplementary figure 4A). To better evaluate the impact of STING knockout on T cell polarization we developed a 16-colors flow cytometry panel dedicated to T cell subpopulation analysis (supplementary figure 4B and C). This showed that while CD103^+^ Treg proportion was decreased, the frequency of T-bet^+^ CD4^+^ and T-bet^+^ CD8^+^ T lymphocytes increased among total immune cells in STING-Ko tumors (irrespectively of their Ki67, CD39, PD1 and CD69 expression). This suggested that STING knockout favored the enrichment of IFNγ producing type-1 helper (Th1) CD4^+^ T lymphocytes and CD8^+^ T cell polarization toward the acquisition of cytotoxic functions^31^ (figure 5D). These differences were also confirmed by analyzing the proportion of CD103^+^ Tregs among CD4^+^ T cells and of T-bet^+^ cells among CD4^+^ and CD8^+^ lymphocytes suggesting a qualitative rather than quantitative modification in T cell polarization in STING-Ko M8 tumors (figure 5E).

By analyzing the myeloid compartment of these tumors, we also noticed that although the proportion of total neutrophils was unchanged, the frequency of the SiglecF^+^ sub-population, which contains CD80^+^MHC-II^+^ double positive cells, was significantly decreased among CD45^+^ immune cells (figure 5F and supplementary figure 4D).

Finally, the proportion of CD103^+^Tregs among CD4^+^T cells was positively correlated with the frequency of SigecF^+^ neutrophils among immune cells, while CD103^+^ Tregs and SiglecF^-^ neutrophils were not associated together (figure 5G). Altogether, these observations showed an unexpected impact of STING knockout in M8 cells highlighting a qualitative change of CD4^+^ and CD8^+^ T lymphocyte pools in favor of T-bet^+^ Th1 and cytotoxic-like cells. They also suggest a functional link between SiglecF^+^ neutrophils and CD103^+^ Treg in M8 tumors.

## Discussion

In this study, we present and characterize two transplantable tumor models derived from the KP mouse. These two models are particularly interesting as they form tumors showing different immune infiltration. While the Clem2 tumors are highly infiltrated by CD8^+^ T lymphocytes and can be cured by anti-PD1 treatment as single agent, the M8 cells generate lesions highly infiltrated by SiglecF^+^ neutrophils and resistant to immunotherapy. Furthermore, the Clem2 cells line is characterized by a high expression of STING and showed basal STING signaling while M8 cancer cells demonstrated low STING and cGAS expression without any clear marker of endogenous STING/IRF3 signaling *in vitro*. Hence, based on these observations and on our bioinformatics analyses showing that STING expression associates with a longer survival in *KRAS* mutant lung cancer patients, we were expecting to generate consistent observations regarding pre-existing literature and showing that STING signaling counteracts lung tumor growth by favoring the establishment of a cytotoxic T-cell anti-tumor immunity^20–22^.

However, our results depict a more complex picture of STING signaling in lung cancer cells suggesting a functional duality. Indeed, our major observation was that in the STING-high Clem2 model, STING knockout impairs subcutaneous tumor formation and that it decreases lung colonization after tail vein injection of the STING-low M8 cells. We are currently performing experiments to identify downstream molecular mechanisms that could help understanding how STING might promote M8 and Clem2 cell transplantation. Our current hypothesis is that STING-mediated CCL5 secretion helps tumor formation *in vivo*^25,32^. This critical involvement of STING signaling in our lung cancer cell lines goes in line with a previous publication showing that chromosomal instability in MDA-MB-231 cells drives STING dependent non-canonical NF-kB activation that is required for metastasis formation^33^. While STING agonist-based therapeutic approaches are currently under investigation in clinical settings in lung cancer^19^, our observations raise important questions on the impact that these new therapeutics could have on cancer cell aggressiveness.

We failed to observe a clear impact of STING knockout on Clem2 tumor immune infiltration. This latter observation is consistent with a previous report that showed that cGAS but not STING expression by the cancer cells can contribute to anti-tumor immunogenicity through a transfer of 2’3’cGAMP to the immune cells^34^. However, we detected a trend toward a faster tumor growth and the response to anti-PD1 immunotherapy was delayed, when transplanting an important number of cancer cells to overcome the deleterious impact of STING knockout on tumor formation. Although we did not treat mice with genotoxic agents to activate STING and analyze its impact on the tumor immune environment, we are not expecting drastic differences, as Clem2 cells did not demonstrate strong induction of ISGs *in vitro*. Furthermore, when compared to TLR3 triggering, STING activation with optimal doses of its specific ligand 2’3’cGAMP led to modest transcription of *Infb* in Clem2 and M8 cancer cell lines and also in MEFs. This observation questions the relevance of STING signaling in cancer cells as potent inducer of interferon, especially in cells showing a general inhibition of ISGs induction capacity such as Clem2. Indeed, these cells showed a low ability to upregulate ISG expression upon both STING and TLR3 triggering when compared to M8 cells and MEF. These results rather suggested that Clem2 cells could have acquired the ability to circumvent IRF3 target genes over expression. STING has been shown to drive non-canonical NF-kB signaling in dendritic cells that repress *Ifnb* expression^35^. Hence, it is possible that in Clem2 cells, a constitutive activation of non-canonical NF-kB signaling represses ISG induction, and STING itself might be involved.

Contrasting to Clem2, M8 cells, characterized by low levels of both STING and cGAS, showed a clear induction of *Ccl5* and *Cxcl10* upon STING triggering. Surprisingly, STING-Ko M8 tumors displayed a slightly better response to anti-PD1 treatment as single agent with a lower infiltration by SiglecF^+^ neutrophils and CD103^+^ Treg accompanied by a high proportion of T-bet^+^ lymphocytes. We previously demonstrated that neutrophil infiltration leads to the exclusion of other immune effector cells, especially CD8 T cells, from the tumor mass in about 20% of NSCLC patients and renders autochthonous KP tumors insensitive to anti-PD1 immunotherapy^36^. Interestingly, tumors from this mouse model were shown to be highly infiltrated by a sub-population of tumor promoting neutrophils expressing SiglecF^28,29^. It is now generally accepted that tumor-associated neutrophils (TANs) participate in resistance to PD/PDL1 inhibition in multiple models of lung cancer and other cancers^9,37–40^. Furthermore, we have shown, using different TAN depletion methods^41^, that these cells can drive immunosuppressive Treg accumulation in the tumor environment^36,42^. Hence, our observations suggest that STING contributes to SiglecF^+^ neutrophil accumulation in tumors, favoring CD103^+^ Treg recruitment and/or amplification that in turn could inhibit T-bet^+^ T lymphocyte enrichment and early anti-PD1 response. Even if the validation of such a scenario requires the identification of the mechanisms through which STING expression in M8 cells favors the accumulation of SiglecF^+^ neutrophils, we trust that STING signaling plasticity in tumor cells warrants further attention.

Finally, further analyses are ongoing to determine if high-STING expression could be a marker of cancer cell dependency to CCL5/CCR5 signal transduction and non-canonical NF-kB activation. Following the observation that STING low expression by the cancer cells can contribute to tumor immunosuppression through enhanced SiglecF^+^ neutrophil and CD103^+^Treg accumulation in basal condition, it will be interesting to trigger its activation with genotoxic agents to see if, in this situation, low STING expression can drive the establishment of an anti-tumor immune microenvironment as proposed by others. This manuscript presents a complex picture of STING signaling in cancer cells that will complement already established knowledge linking extranuclear DNA sensing in the cancer cell to anti-PD1 immunotherapy success.

## Methods

### Cell lines origin, culture and treatment in vitro

SV2, M8, CLEM2 and CLEM42 were generated from autochthonous tumors obtained in the *Kras^lox-Stop-LoxG12D/wt^;p53^fl/fl^* (KP) mouse model of lung cancer. Primary lung tumor formation was initiated by intra-tracheal administration of a lentivirus coding for the Cre-recombinase to induce *Kras^G12D^* expression and completely abrogate *p53* translation in lung epithelial cells. At end-point (6-8 months post tumor-initiation), mice were sacrificed and one tumor per animal was dissociated and put in culture in DMEM medium supplemented by 10% fetal calf serum (FCS) and penicillin plus streptomycin. After 15 passages cells were showing stable proliferation rate and were frozen. All of these cell lines had been validated for their ability to form tumors in the lung of C57BL/6J syngeneic mice after tail vein injection of 50 000 cells.

STING or TLR3 triggering was achieved using 2’3’cGAMP (Ref: SML 1229) or Polyinosinic-polycytidylic acid potassium salt (poly (I:C), (Ref: P9582) purchased from Sigma-Aldrich. 2’3’cGAMP and pI:C were formulated in lipofectamine at a final concentration of 10μM and 2μg/ml respectively.

### Generating ΔN-IRF3 and STING-KO cell lines

To generate ΔN-IRF3 inducible models, CLEM2, M8 and MEF cells were transduced with a construct that allows doxycycline inducible ΔN-IRF3 expression and resistance to blasticidin. Briefly, a truncated form of the mouse IRF3 lacking the DNA binding motif (133 amino acids at the N-terminal position) was obtained by amplifying the mouse Irf3 cDNA using the fallowing primers; forward: 5’ ATG TCC CAG GAA AAC CTA and revers 5’: TTA TCG TCA TCG TCT TTG TAA TCG ATA TTT CCA GTG GC. This strategy allowed the addition of a FLAG on the C-Terminal part of the mutant protein. Then this sequence was cloned downstream to the tetracycline response element (TRE) in a PJK-rTTa-IRES-BasticinideR Lentiviral vector. After transduction, cells were constantly kept under blasticidin (100μg/ml final, Ref: ant-Bl-05, Invivogen) to maintain selection pressure; to induce ΔN-IRF3, doxycycline (SIGMA-Aldrich, Ref: D9891) was used at 1μM. All the cell lines were cultured under 5% CO2 at 37° C in DMEM media supplemented with 10% FCS and 1% gentamicin. Cells were frequently tested to ensure that they were not infected by Mycoplasma. STING knockout cell lines were generated using GFP and RFP lentiviral vectors (Addgene, ref 57818 and ref 57819) in which the following guild-RNA were cloned: GFP vector: fw 5’ CACC GTCCAAGTTCGTGCGAGGCT and rv: 5’ AAAC AGCCTCGCACGAACTTGGAC; RFP vector: fw: 5’ CACCGTAGAGAGCTTTGGGGCCTC and rv: 5’ AAAC GAGGCCCCAAAGCTCTCTAC. Cell lines were then transduced with both GFP and RFT vectors containing guild-RNA (STING-Ko) of without guild-RNA (STING-wt). Green and red double positive cells were sorted by flow cytometry to generate polyclonal M8 and Clem2 STING-Ko and STING-wt cell lines.

### Real Time PCR

Total cellular RNA was extracted with TRI reagent (MERK/SIGMA-Aldrich Ref:T9424) accordingly to manufacturer instructions. 1μg of RNA was retrotranscribed in cDNA using PrimSTAR superscript III (ThermoFisher Ref:1808005). 10 ng of cDNA were analysed by qPCR in duplicates using ONEGreen FAST qPCR Premix (OZYME Ref: OZYA008-40/OZYA008-200XL). The qPCR reactions were run on Light Cycler 480 II (Roche) instrument. B-actin or GADPH were used as housekeeping gene for normalization.

Primer sequences:

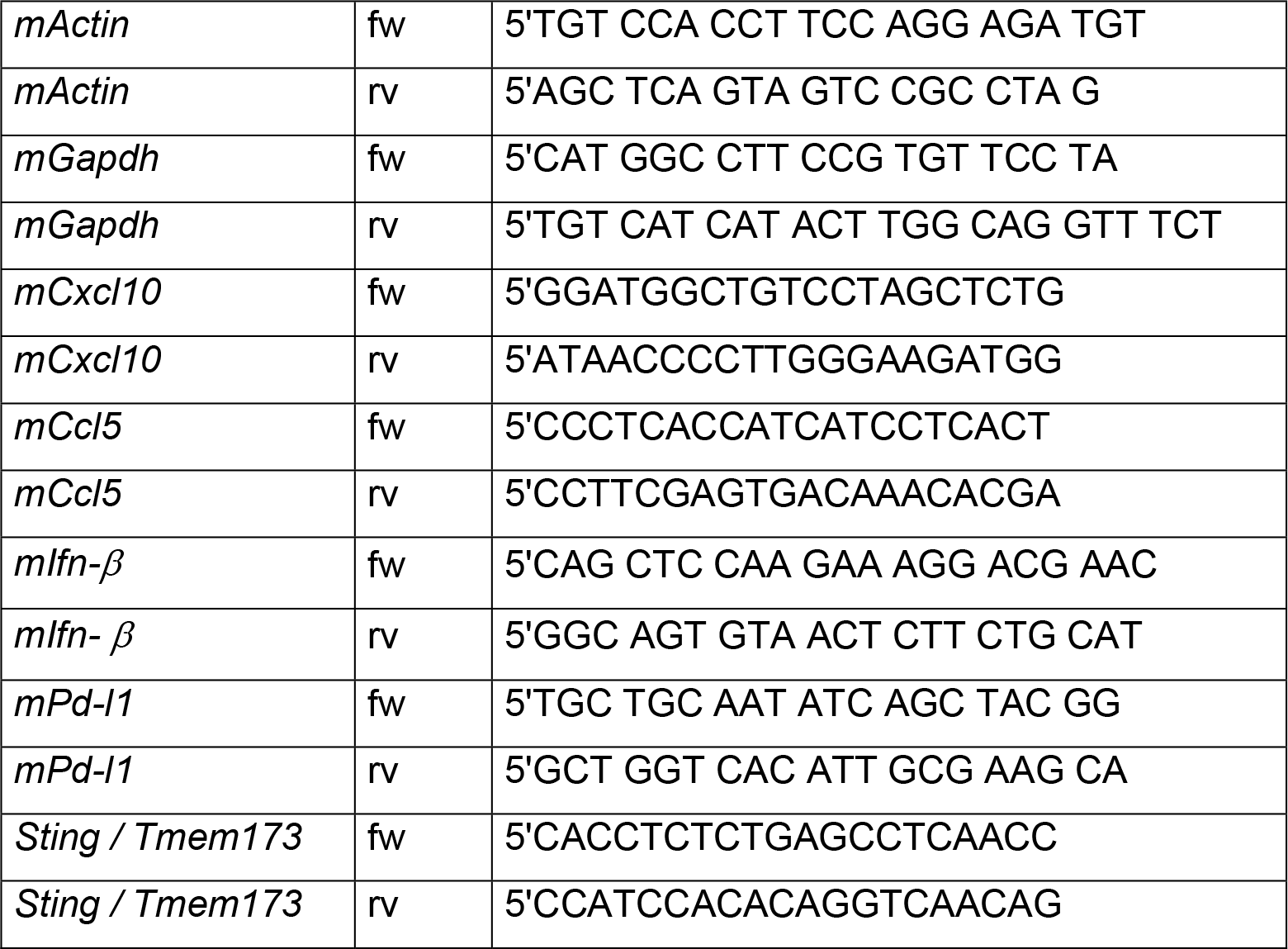

### Western Blotting

Cells were lysed in Ripa Buffer (50 mM Tris HCl pH 8, 150 mM NaCl, 5 mM EDTA, 0,5% DOC Na, 0,1% SDS, 1% TritonX100, 1% phosphatase inhibitor, 2% proteinase inhibitor, 2,5% SDS). 50 or 135 □g of total proteins were loaded on 10% polyacrylamide gels and electro-transferred on nitrocellulose membranes. Membranes were blocked in TBS with 0,1% of Tween 20 (TBS-Tween) and 5% of milk (1-2h RT), then incubated overnight at 4° with the primary antibodies: STING (Protein tech Ref: 19851-1-AP, Polyclonal), HSC70 (Santa Cruz Biotechnology, (B-6) Ref: sc-7298), ACTIN (Cell signalling, Clone13E5, Ref: 4970), E-CADHERIN (Cell signalling, Clone24E10, Ref: 3195), VIMENTIN (Cell signalling, CloneD21H3, Ref: 5741), C-GAS (Cell signalling, CloneD3080, Ref: 316595), TBK1(Cell signalling, CloneD1B4, Ref: 3504S), IRF3 (BioLegend, Clone12A4A35, Ref: 655701) (all of them diluted 1/1000 with the exception of TBK1 that has been diluted 1:500). The membranes were than washed three times in TBS-Tween and incubated with the secondary antibody IRDye 680RD anti-rabbit IgG (Ref: 925-68071) or IRDye 800 CW anti-mouse IgG (Ref: 925-32210) were purchased from LI-COR and used a dilution 1/10 000. Membranes have been left to dry and revealed with LI-COR ODYSSEI FC imaging system.

### Immunodetection of cytosolic double-stranded DNA (dsDNA) in Clem2 and M8 cells

Based on the protocol by Spada et al^43^, Clem2 and M8 cells grown in 12 mm diameter coverslips were fixed with 4% PFA and permeabilized for 7min. After blocking nonspecific sites, cells were incubated with an anti-dsDNA antibody (ab27156, Abcam), diluted to 1:1000 in 1% BSA/PBST, at 4 °C overnight. After three washes in 1X PBS, cells were incubated with a goat anti-mouse IgG Alexa Fluor^®^ 488 at room temperature for 1h. After three washes in 1X PBS, coverslips were mounted on a slide using one drop of Vectashield^®^ antifade mounting medium with DAPI (H-1200, Vector Laboratories). Images were acquired at 40x magnification using a Zeiss^®^ Apotome.2 (AxioCam MRm) microscope.

### Murine tumor models and treatments

8 weeks-old male C57BL/6J mice were purchased from Charles River. For tumor growth mice were kept in specific pathogen-free conditions in the animal facilities of the Institut de Recherche sur le Cancer de Montpellier. All procedures for animal handling and experiments were approved by the ethics committee of the local animal facility (‘‘ComEth’’) and the institutional review board, under the authority of the regional ethics committee for animal experimentation.

To generate subcutaneous tumor models, Clem2 cells and M8 cells suspended in 100μl PBS were injected subcutaneously (s.c.) into 8 weeks-old male C57BL/6J. Tumor growth was monitored three times per week with a caliper and mice were killed when the tumor reached a volume of 800 mm3. Tumor volume was calculated with the following formula: length x width x width/2. Orthotopic transplantation were performed using 5×10^4^ M8-wt and Clem2-wt cells suspended in 100μl PBS and injected trough tail vain injection.

For assessment of the therapeutic effect of anti-PD-1 mAb, M8- and Clem2-grafted mice received intraperitoneal (i.p.) injections of 150 μg mAb in a volume of 100μl, twice a week during 4 weeks. The anti-PD1 (clone RMP1-14, Ref: BE0146) was purchased from BioXcell and treatment started at day 10 post-transplantation when the average tumor volume was of 40-70 mm3. Tumor volume or percent of survival versus time were plotted on graph using GraphPad Prism software.

### Sample preparation for flow cytometry

200μl of blood was recovered in heparine tubes and red blood cells were eliminated by adding 2 mL ACK lysing buffer. White blood cells were recovered by centrifugation, washed with PBS and suspended in flow cytometry buffer (2% FCS, 0.5 M EDTA, 0.02% NaN3 in PBS) for staining and flow cytometry analysis. The weight of Isolated tumors or lungs was systematically measured, 350mg of tissue was minced to generate 2 mm^3^ fragments and placed in RPMI1640 containing 200μg/ml of collagenase type IV (ref: C4-BIOC,Sigma-Aldrich) and 50U/ml of DNase I (11284932001, Roche) then incubated using the a gentleMACS dissociator (milteniy). After dissociation, single cell suspensions were passed through 70 μm filters with (Falcon; Cell Strainer) and centrifuged. Cell pellets from tumor single cell suspensions or blood were resuspended in 100μl of flow cytometry buffer per 20mg of tissue or 100μl of blood in flow cytometry buffer for antibody staining.

### Flow cytometry antibody staining and true-count

For each samples we always used a volume of cell suspension equivalent to 30mg of tissue or 100μl of blood. Cells were incubated with mouse Fc-blocking reagent (Miltenyi Biotec) plus Viakrom-808 (Bekman Culter) for 20 min at 4° C in flow cytometry buffer, then the appropriate antibody cocktail was added for 20min at 4°C. Cells were then fixed with 1% PFA in PBS of in Foxp3 staining buffer kit (ebioscience) and stored until acquisition with the Cytoflex LX-. Data were analyzed with Flowjo 10.7 software (FlowJo LLC ©). For true-count flow cytotrey, 20.103 absolute count beads (biolegend) were added to the single cell suspension before cytometry acquisition

Antibody list:

|  |  |  |  |  |  |
| --- | --- | --- | --- | --- | --- |
| CD3 | 100220 | Biolegend | CD274 (PD-L1) PE/Dazzle™ 594 | 124324 | Biolegend |
| CD3-PE | 130-121-133 | Milteniy | CD279 (PD-1) PE/Dazzle™ 594 | 135228 | Biolegend |
| CD4 Biotine | 130-118-696 | Milteniy | CD335 (NKp46) Biotin | 130-112-356 | Milteniy |
| CD4 BUV 456 | 741134 | BD Biosciences | F4/80 Brilliant Violet 605™ | 123133 | Biolegend |
| CD8 BUV 661 | 750023 | BD Biosciences | F4/80 PE/Cyanine7 | 157308 | Biolegend |
| CD8a-Biotin | 130-118-147 | Milteniy | FcR Blocking Reagent, mouse | 130-092-575 | Milteniy |
| CD11b-FITC | 130-113-234 | Milteniy | FoxP3-APC | 17-5773-82 | eBioscience |
| CD11c Brilliant Violet 421™ | 117330 | Biolegend | I-A/I-E (MCHII) BUV 395 | 743876 | BD Biosciences |
| CD19 Biotin | 130-112-034 | Milteniy | Ki-67 eFluor780 | 47-5698-82 | eBioscience |
| CD39-PerCP-eFluor 710 | 46-0391-82 | eBioscience | Ly6C-PerCP | 130-111-920 | Milteniy |
| CD45 | 103149 | Biolegend | Ly-6G Alexa Fluor® 700 | 127622 | Biolegend |
| CD45R (B220)-FITC | 130-110-845 | Milteniy | Ly-6G Brilliant Violet 605™ | 127639 | Biolegend |
| CD45R/B220 Brilliant Violet 650™ | 103241 | Biolegend | NK-1.1 Alexa Fluor® 700 | 108730 | Biolegend |
| CD68 Alexa Fluor® 700 | 137026 | Biolegend | Precision Count Beads | 424902 | Biolegend |
| CD69 Brilliant Violet 650™ | 104541 | Biolegend | Siglec-F-PE | 130-112-174 | Milteniy |
| CD103 Brilliant Violet 510™ | 121423 | Biolegend | T-bet Brilliant Violet 421™ | 644816 | Biolegend |
| CD103 | 121406 | Biolegend |  |  |  |

### Statistics

For all in vitro experiments Statistical analysis was performed using Prism 8 software. All results are represented as mean ± SEM if not stated otherwise. Comparisons between groups were made as stated in the figure legends. Statistical significance is indicated as *p < 0.05, **p < 0.01, ***p < 0.001, ****p < 0.0001, or ns (not significant), using Mann-Whitney tests, where not indicated otherwise.

Bioinformatics analysis were performed based on identical R packages.In the TCGA LUAD database, we identified 198 TCGA patients for which mutation status is known and split by median expression of STING. Among the 198, 138 are *KRAS* WT and 60 are *KRAS* MUT.For CCLE analyses, the geneset collection used for GSEA^26^ is the HALLMARK collection from mSigDB^27^.

## Author contributions

LG and CU designed and performed experiments and contributed in interpreting the data and writing the manuscript. JC performed cytoplasmic DNA staining. NZ performed bioinformatics analyses of the TCGA and CCLE datasets. EM provided the mouse KP tumor and MEF cell lines and JF produced ΔN-IRF3 and CrispR vectors under his supervision. NB contributed in data interpretation and edited the manuscript. JF initiated and supervised the study and wrote the manuscript.

## Abbreviations

CCLE: cancer cell line encyclopedia
cGAS: cyclic GMP-AMP synthase
2’3’-cGAMP: cyclic Guanosine(2’,5’)phosphate-Adenosine(3’,5’)phosphate
IFN: interferon
ISG: Interferon-stimulated genes
IRF3: Interferon regulatory factor-3
MEF: mouse embryonic fibroblasts
NSCLC: non-small cell lung cancer
PD1: programmed cell death receptor-1
STING: Stimulator of Interferon Gene
TBK1: Tank-binding kinase-1
TCGA: the cancer genome atlas

