## Supplemental figures for "Low STING expression in a transplantable Kras^G12D^/P53^ko^ lung cancer model contributes to SiglecF^+^ neutrophil and CD103^+^Treg accumulation in tumors"

**Gros et al:**

### Supplementary figure legends

#### Supplementary figure 1: Immunophenotyping of Clem2 and M8 tumors

A) Spleen weight (left) and number of $CD45^{+}$ leukocytes (right) per ml of blood of age matched naïve mice (n=5 and n=3) and at end-point after tail vein transplantation of Clem2 (n=3) or M8 (n=6 and n=5) cells. * $p<0.05$, Statistics were obtained using Mann Whitney test.

B) Representative flow-cytometry plots depicting the gating strategy used in Clem2 (left) and M8 (right) sc-tumors to quantify immune cells per mg of tissue. First gating level shows identification of absolute count beads.

#### Supplementary figure 2: Exploring STING and IRF3 signaling in lung cancer cell lines and MEF in vitro

A) Western blot showing endogenous-IRF3 (endo) and ΔN-IRF3, STING and HSC70 expression in MEF, M8 and Clem2 cells in control condition or following 72 hours of doxycycline exposure (Dox).

B) Histograms shows relative mRNA expression of the indicated gene in MEF, Clem2 and M8 cells exposed medium containing lipofectamine or lipofectamine plus 2 µg/ml of pI:C for four hours. DCT were calculated based on *Gapdh* expression. Results from one representative experiment over three. Numbers indicate fold change from respective control without pI:C.

C) Histograms depict the results from proliferation (left) wound healing (middle) and colony formation assays on Clem2-wt, Clem2-Ko, M8-wt and M8-Ko cells.

B-C) * $p<0.05$, ** $p<0.01$, ***$p<0,001$ and **** $p<0.0001$ Statistics were obtained using Student-T test.

**Supplementary figure 3: STING knockout reduces M8 cancer cells ability to colonize the mouse lung**

A) Histogram shows the quantity of $CD45^{+}$ cells in Clem2-wt (n=5) and Clem2-Ko (n=5) sc-tumors at end-point from true-count flow cytometry.

B-C) Histograms represents the percentage of CD4+ T cells (left) CD8+ T cells (right) (A) and the ratio CD8+ T cells over Treg (B) in Clem2-wt (n=5) and Clem2-Ko (n=7) sc-tumors at end-point.

D) Curves represent the growth of M8-wt (black n=8) and M8-Ko (blue n=8) sc-tumors.

E) Kaplan–Meier curves showing mice overall survival of mice after tail vein injection of M8-wt (black n=7) and M8-Ko (blue n=8) cells. Statistic was obtained using Wilcoxon test, $*p<0.05$

**Supplementary figure 4: Immunophenotyping of M8 STING wild type and knockout sc-tumors**

A) Expression of CD39 and Ki67 on $CD8^{+}$ T cells, T-$bet^{-}$, T-$bet^{+}$, $Foxp3^{+}CD103^{-}$ and $Foxp3^{+}CD103^{+}$ $CD4^{+}$ T cells in an M8 sc-tumor.

B-C) Representative flow cytometry plots showing the gating strategy used to identify T-$bet^{-}$, T-$bet^{+}CD8^{+}$ T cells, $Foxp3^{-}$ T-$bet^{-}$ and T-$bet^{+}$ $CD4^{+}$ T and $Foxp3^{+}CD103^{-}$ and $CD103^{+}$ Tregs in M8 sc-tumors (B) while simultaneously monitoring their expression of PD1, CD39, Ki67 and CD69 (C).

D-E) Representative flow cytometry plots showing the gating strategy used to identify $SiglecF^{+}$ neutrophils among immune cells (D) and to monitor their expression of CD80 and MHCII (E) in M8 sc-tumors.

**Supplementary figure 1:**
**Immunophenotyping of Clem2 and M8 tumors**

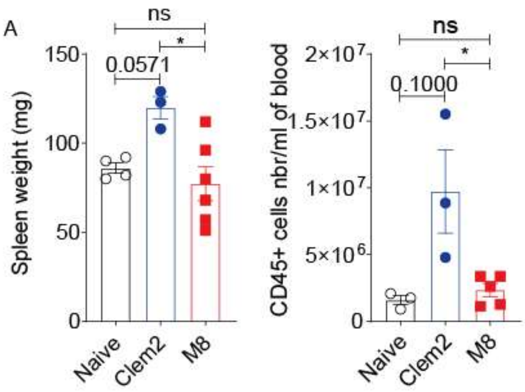


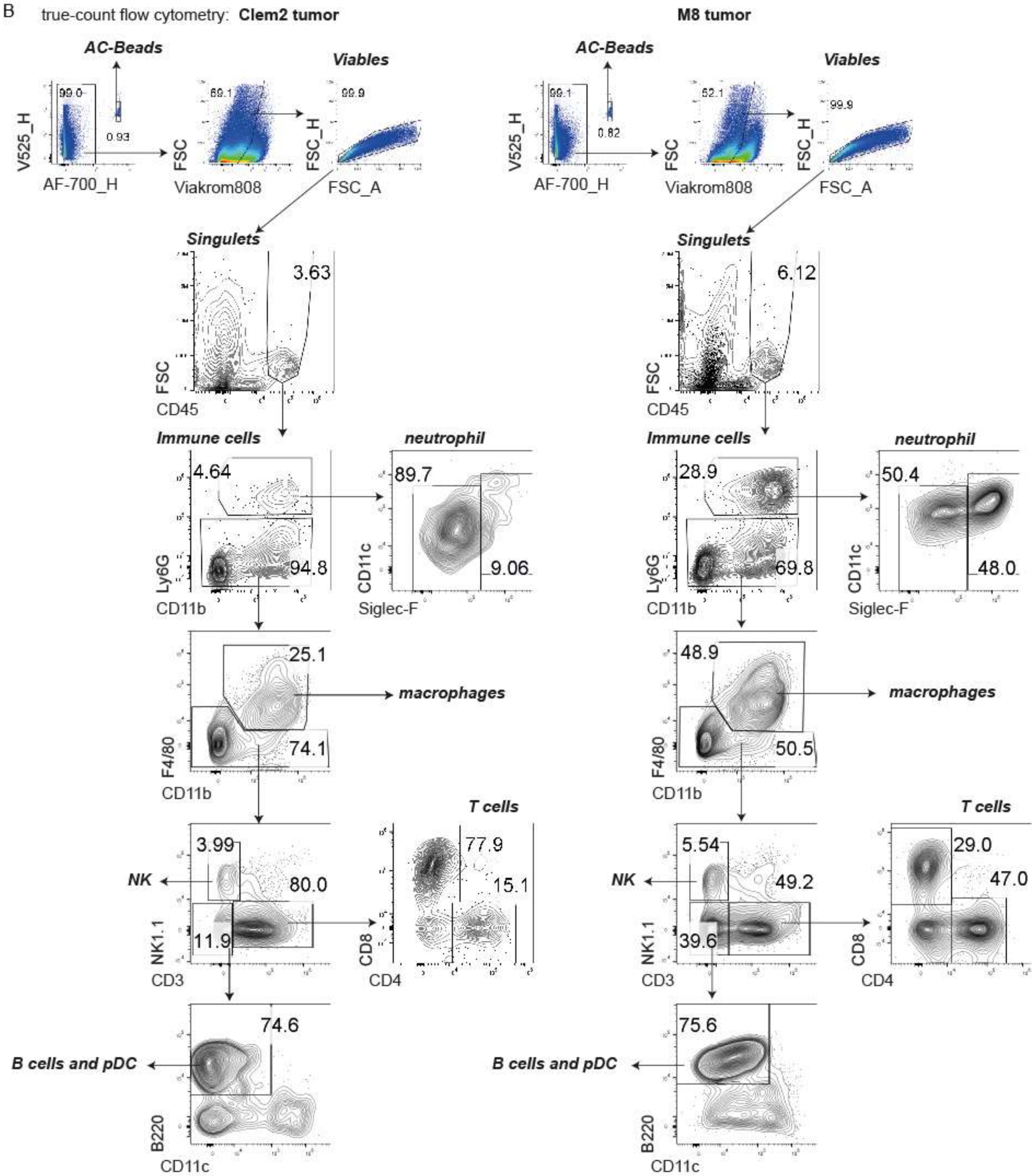

**Supplementary figure 2: Exploring STING and IRF3 signaling in lung cancer cell lines and MEF in vitro**

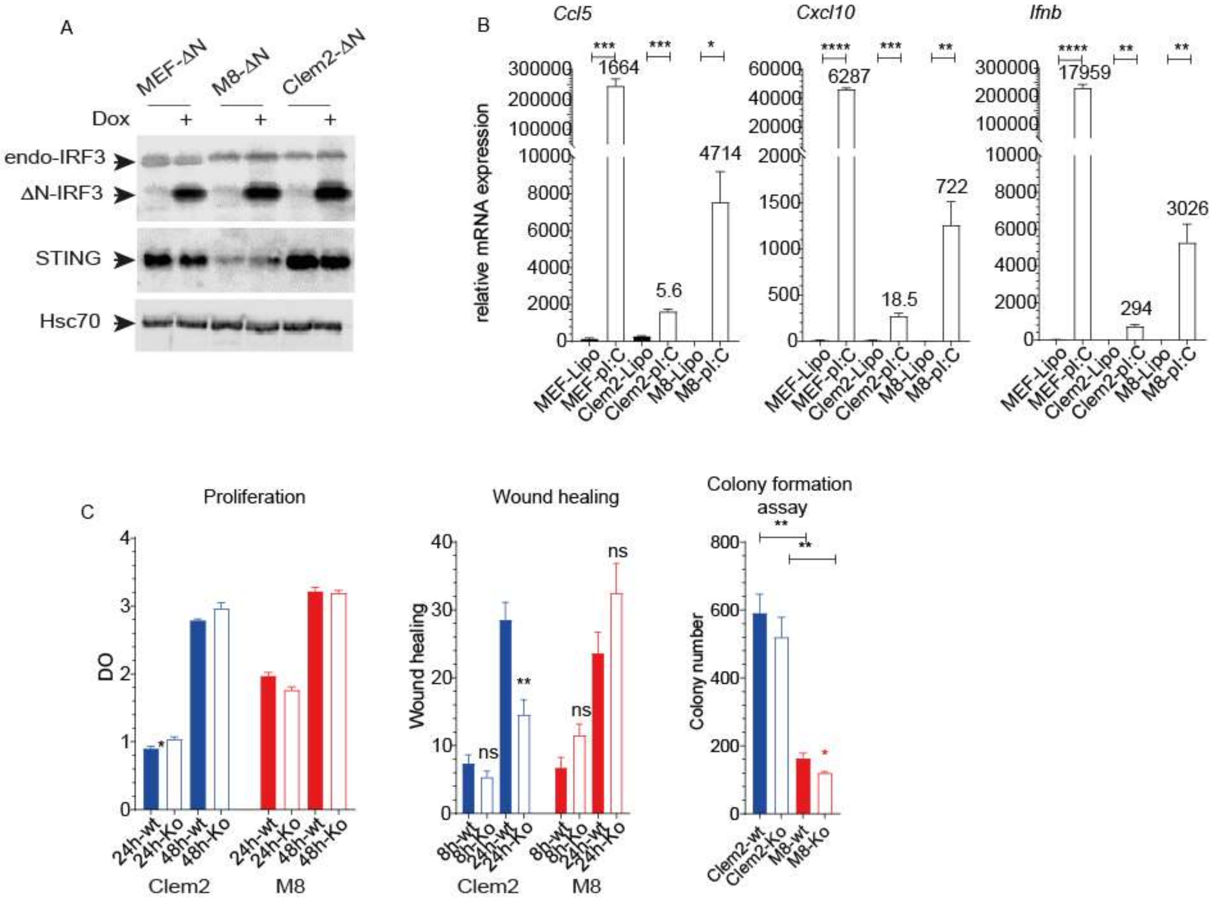

**Supplementary figure 3: STING knockout reduces M8 cancer cells ability to colonize the mouse lung**

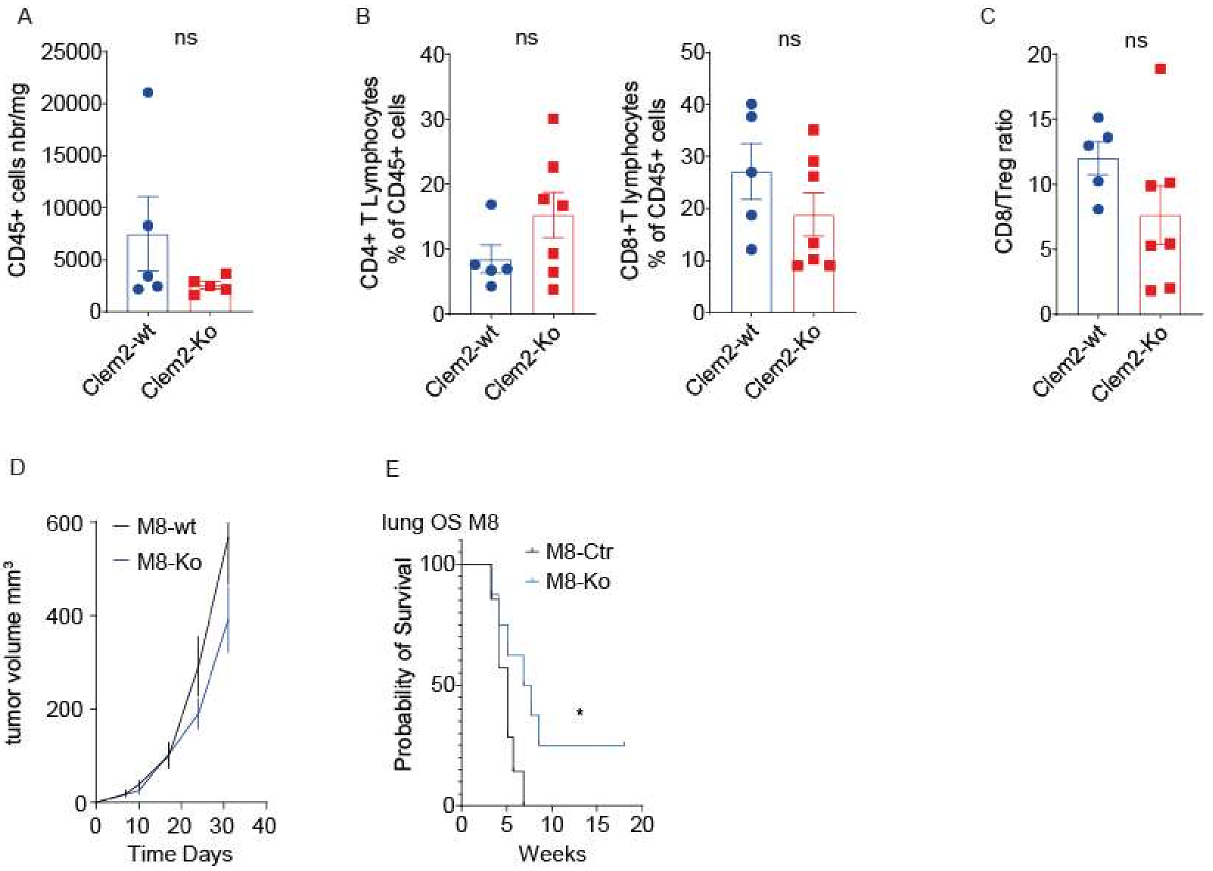

**Supplementary figure 4: Immunophenotyping of M8 STING wild type and knockout sc-tumors**

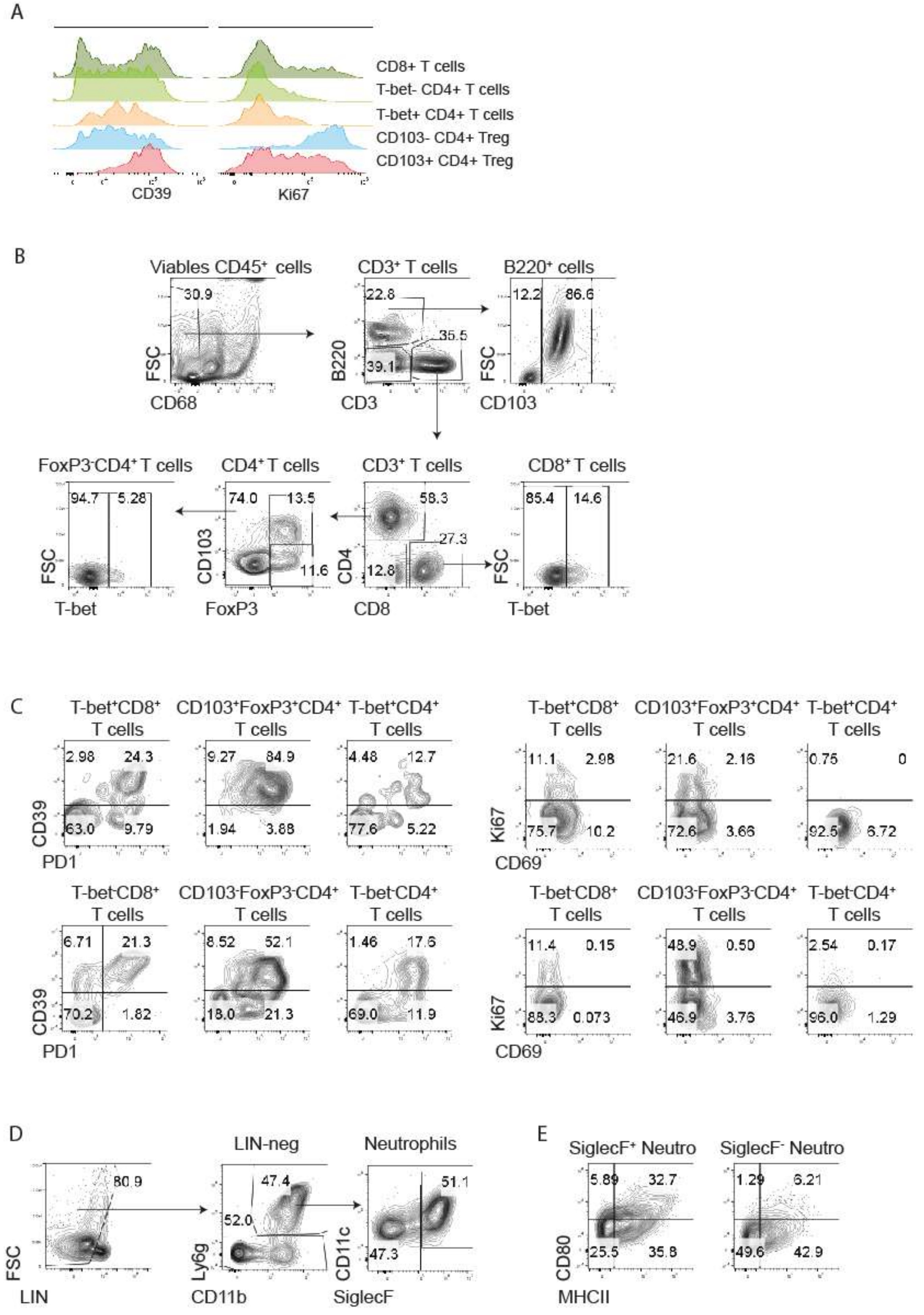
